# Dual Targeting of *BRAF^V600E^* and Ferroptosis Results in Synergistic Anticancer Activity via Iron Overload and Enhanced Oxidative Stress

**DOI:** 10.1101/2025.06.25.661418

**Authors:** Jiangnan Hu, Chandrayee Ghosh, Tejinder P. Khaket, Zhongyue Yang, Yasmine Tabdili, Eden D. Alamaw, Myriem Boufraqech, Scott J. Dixon, Electron Kebebew

**Affiliations:** Department of Surgery and Stanford Cancer Institute, Stanford University, Stanford, CA, USA; University of Arizona College of Medicine, Tucson, AZ, USA; Department of Comparative Medicine, Stanford University, CA, USA; Endocrine Oncology Branch, Center for Cancer Research, National Cancer Institute, NIH, MD, USA; Department of Biology, Stanford University, Stanford, CA, USA

**Keywords:** Anaplastic thyroid cancer, BRAF mutation, ferroptosis, GPX4, combination therapy

## Abstract

**Purpose:** While combination BRAF and MEK inhibitor treatment in *BRAF^V600E^*-mutant cancers results in a response, treatment resistance and toxicity are common. Ferroptosis is an iron-dependent form of non-apoptotic cell death. BRAF inhibition has been associated with increased sensitivity to ferroptosis that is dependent on Glutathione Peroxidase 4 (GPX4).

**Experimental Design:** *In vitro*, *ex vivo*, and *in vivo* models of anaplastic thyroid cancer (ATC) were used to evaluate the anticancer activity of combination BRAF inhibition and ferroptosis induction.

**Results:** Targeting key regulators of ferroptosis—GPX4, using RSL3 and ML162, and system X_c_^−^, using erastin—induced significant cell death in all ATC cell lines. Combination of dabrafenib and RSL3 synergistically increased cell death in *BRAF^V600E^*-mutant ATC cells, and significantly inhibited cellular migration and colony formation. Mechanistically, lipid peroxidation, reactive oxygen species levels, and intracellular Fe^2+^ increased significantly with combination treatment compared with each agent alone. Analysis of cell membrane iron importers and exporters showed significantly lower expression of ferroportin-1 (an iron exporter), suggesting the synergistic anticancer activity was due to increased iron accumulation and oxidative stress, leading to enhanced ferroptotic cell death. *BRAF^V600E^*-mutant ATC cell spheroids showed synergistic cell death with dabrafenib and RSL3 treatment. *In vivo*, combination dabrafenib and ferroptosis induction (by targeting GPX4 using C18, and system X_c_^−^ with IKE) significantly inhibited tumor growth in an orthotopic ATC mouse model. Additionally, dabrafenib-resistant *BRAF^V600E^*-mutant ATC cells were more sensitive to ferroptosis induction than parental cells.

**Conclusions:** Dual targeting of *BRAF^V600E^* and ferroptosis results in synergistic anticancer activity and overcomes resistance to BRAF inhibition.

## INTRODUCTION

The *BRAF* oncogene is commonly mutated in multiple types of cancers (thyroid, colon, rectum, lung, ovary, and brain) (1). The most frequent *BRAF* mutation is *V600E*; it has been successfully targeted in the clinic using BRAF inhibitor, however, acquired drug resistance is common and leads to a significant relapse in almost all cases (2). Combination therapy with a BRAF inhibitor and a MEK inhibitor was initially investigated in *BRAF^V600E^*-mutant metastatic melanoma and demonstrated greater effectiveness than BRAF inhibition alone (3). Although this combination treatment results in a clinical response, most patients develop resistance and treatment-related toxicities as reported recently (4). Moreover, because both BRAF and MEK inhibitors act largely on the same pathway, the clinical response is often not durable (5). Anaplastic thyroid cancer (ATC) has one of the highest mortality rates of all human malignancies, and approximately 50% of patients with ATC have the *BRAF^V600E^* mutation (6). While combination BRAF and MEK inhibitor treatment is the first-line treatment in patients with *BRAF^V600E^*-mutant ATC, the response to treatment is also not durable. Therefore, new treatments that result in a significant and durable response are needed in *BRAF^V600E^*-mutant cancers.

Ferroptosis is a non-apoptotic form of regulated cell death that is dependent on intracellular iron and is dysregulated in cancer (7). Thus, targeting ferroptosis for cancer therapy has been actively investigated in recent years. Cancer cells that can tolerate BRAF inhibitors and other treatments can acquire a mesenchymal cell state that is more dependent for survival on glutathione peroxidase 4 (GPX4, a gatekeeper of ferroptosis that protects against ferroptosis), and they present increased sensitivity to ferroptosis induction (8). Additionally, BRAF inhibition can also induce dedifferentiation, which increases sensitivity to ferroptosis in *BRAF^V600E^*-mutant melanoma cells (9). Thus, we hypothesized that ferroptosis induction can have anticancer activity in *BRAF^V600E^*-mutant cancer and that combination ferroptosis induction and BRAF inhibition has synergistic anticancer activity due to targeting dual prosurvival cancer pathways. Thus, in the present study, we evaluated the anticancer activity of BRAF inhibition and ferroptosis induction in *BRAF^V600E^*-mutant and wild type ATC using *in vitro*, *ex vivo*, and *in vivo* models of ATC.

## RESULTS

### Synergistic anticancer activity of dabrafenib and RSL3 treatment in *BRAF^V600E^*-mutant ATC cells

As expected, treatment with dabrafenib resulted in a dose-dependent inhibition of cell proliferation in *BRAF^V600E^*-mutant ATC cell lines (8505C and SW1736) (Fig. 1A). Inhibiting system X_c_^−^ (with erastin) and GPX4 (with RSL3 and ML162) resulted in significant loss of cell viability within 48 h in all four ATC cell lines (Fig. 1B–D). Erastin induced greater reduction in cell viability in the *BRAF^V600E^*-mutant cell lines than in the *BRAF^WT^* ATC cell lines (THJ-16T and C643) (Fig. 1B). The extent of cell viability in the RSL3 and ML162 treatment groups was similar across the four ATC cell lines regardless of the *BRAF* mutation status (Fig. 1C, D). GPX4 expression and the reduced glutathione (GSH)/oxidized glutathione (GSSG) ratio were reduced with erastin, and these changes were partially reversed by ferrostatin-1 (a ferroptosis inhibitor), confirming induction of ferroptosis in the ATC cells (Fig. S1A, B).

**Figure 1.**
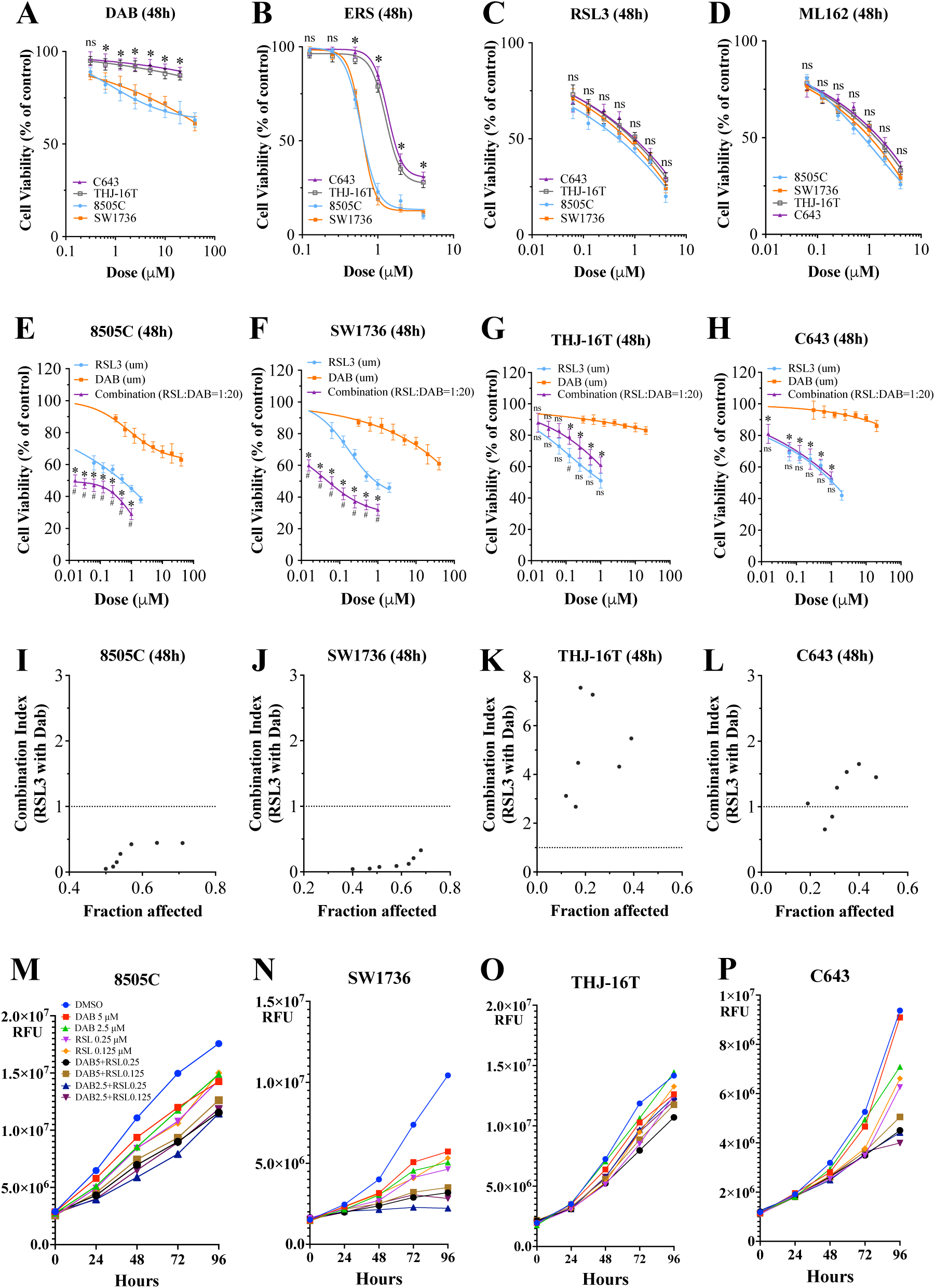
BRAF inhibition and ferroptosis induction in ATC cell lines. (A) Dabrafenib reduced cell viability in a dose-dependent manner in the *BRAF*^V600E^-mutant ATC cell lines (8505C and SW1736). (B) Erastin treatment reduced cell viability in a dose-dependent manner in the ATC cell lines. (C) RSL3 treatment reduced cell viability in a dose-dependent manner in the ATC cell lines. (D) ML162 treatment reduced cell viability in a dose-dependent manner in the ATC cell lines. All data are presented as the mean ± standard deviation (n = 3), *P < 0.05 in the *BRAF*^V600E^-mutant ATC cell lines (8505C and SW1736) compared with the *BRAF^WT^* ATC cell lines (C643 and THJ-16T). (E– H) Combination BRAF and GPX4 inhibition with RSL3. The cell viability curves of the combination treatment were plotted according to the doses of RSL3 used in the treatment. *P < 0.05 in the combination treatment group compared with the dabrafenib-only treatment group at each drug concentration point; ns = nonsignificant. ^#^P < 0.05 in the combination treatment group compared with the RSL3-only treatment group at each drug concentration point; ns = nonsignificant. (I–L) The combination index was calculated using CompuSyn software. A combination index < 1 was considered synergistic. All cell viability data are presented as the mean ± standard deviation (n = 3). ns = nonsignificant. (M–P) The effect of dabrafenib and RSL3 treatment on cellular proliferation. The cells were treated with dabrafenib (2.5, 5 μM), RSL3 (0.125, 0.25 μM), or combinations of the two for up to 96 h. The X-axis represents the elapsed time in hours, and the Y-axis represents the relative fluorescence units. DAB = dabrafenib, ERS = erastin.

Ferroptosis induction with RSL3 caused the greatest amount of reduced cell viability in the ATC cell lines, so we used it to induce ferroptosis for the combination BRAF inhibition studies. Combination RSL3 and dabrafenib treatment had synergistic antiproliferative activity in the 8505C and SW1736 *BRAF^V600E^*-mutant ATC cell lines (Fig. 1E, F). As expected, we did not observe the synergistic effect in the *BRAF^WT^* ATC cells (C643 and THJ-16T; Fig. 1G, H). We used the Chou–Talalay algorithm to assess drug interactions. Dabrafenib and RSL3 exhibited a synergistic effect in the *BRAF^V600E^*-mutant ATC cell lines (Fig. 1I, J), but in the *BRAF^WT^* ATC cell lines, this combination predominantly showed an antagonistic effect (Fig. 1K, L; Table S1). Combination treatment with dabrafenib and RSL3 inhibited cell proliferation significantly more than each agent alone in the *BRAF^V600E^*-mutant ATC cell lines (Fig. 1M, N), but not in the *BRAF^WT^* cell lines (Fig. 1O, P). There were no synergistic effects when combining dabrafenib and erastin treatment in the *BRAF^V600E^*-mutant (Fig. S1C) and *BRAF^WT^* ATC cell lines (Fig. S1D).

Based on the combination index (CI) values derived from the Chou–Talalay method, we used the two concentrations of each drug that produced the most pronounced synergistic effects for colony formation and migration assays (Table S1). We found that combination dabrafenib (5 μM) and RSL3 (0.25 μM) treatment reduced colony formation more than dabrafenib or RSL3 alone in the *BRAF^V600E^-*mutant 8505C and SW1736 ATC cell lines compared with the control (P < 0.05; Fig. 2A, B). In addition, the combination of dabrafenib (5 μM) and RSL3 (0.25 μM) significantly inhibited cellular migration in the *BRAF^V600E^*-mutant ATC cell lines compared with each agent alone (P < 0.01; Fig. 2C, D) following 24 hours of treatment.

**Figure 2.**
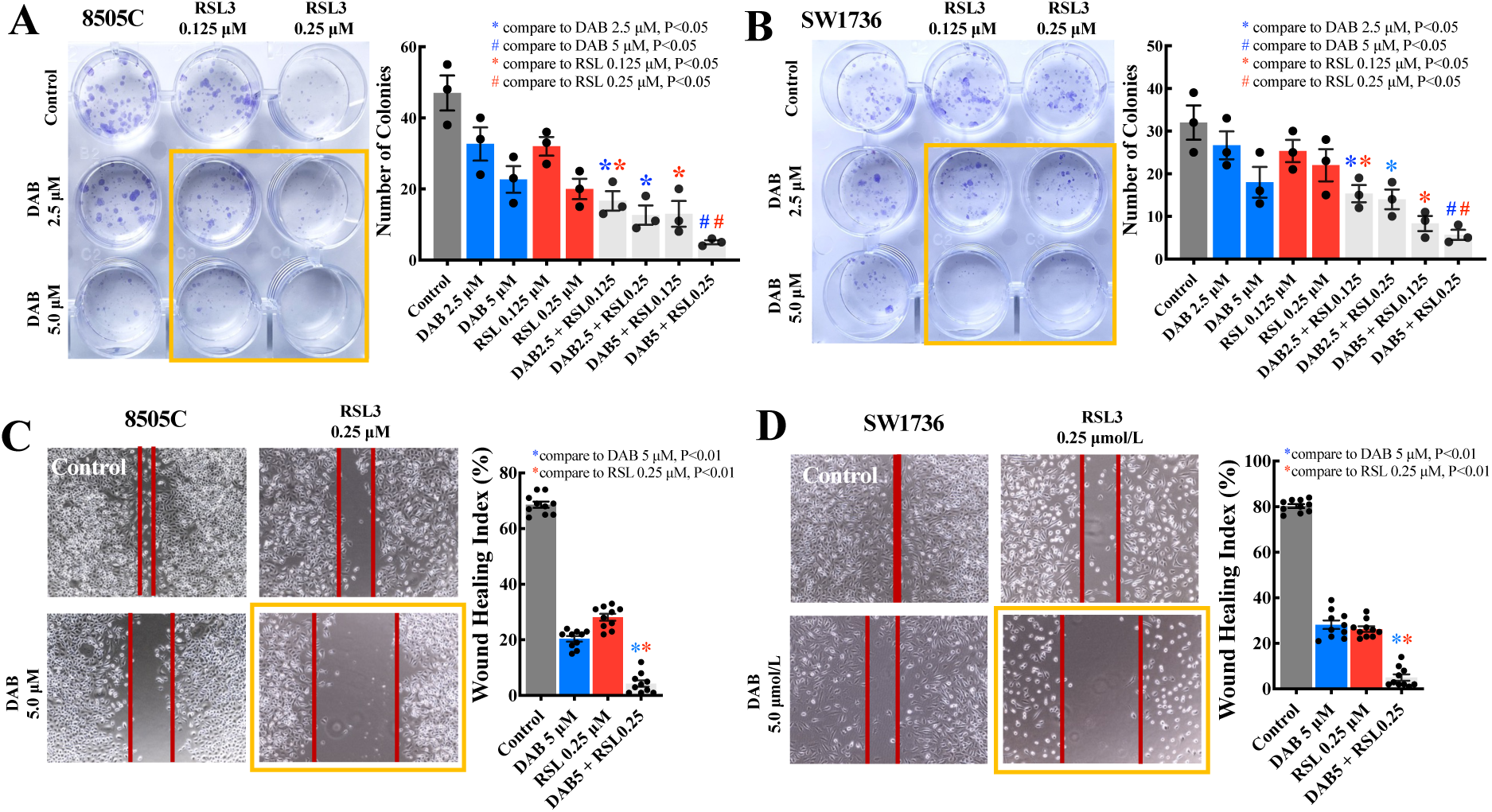
Combination dabrafenib and RSL3 treatment inhibited colony formation and migration in *BRAF^V600E^*-mutant ATC cell lines. (A, B) Combination dabrafenib and RSL3 treatment inhibited colony formation in the 8505C and SW1736 ATC cell lines. The *BRAF^V600E^*-mutant ATC cell lines were incubated with increasing doses of dabrafenib (2.5, 5 μM), RSL3 (0.125, 0.25 μM), or their combinations for 12–14 days. The images are representative of three independent experiments. The histogram presents the mean colony counts of the 8505C and SW1736 ATC cell lines. Combination dabrafenib (5 μM) and RSL3 (0.25 μM) treatment reduced colony formation more than dabrafenib or RSL3 alone in the 8505C and SW1736 ATC cell lines (P < 0.05). (C, D) Combination dabrafenib (5 μM) and RSL3 (0.25 μM) treatment reduced cellular migration more than dabrafenib or RSL3 alone in the 8505C and SW1736 cells ATC cell lines (P < 0.001). All study data are presented as the mean ± the standard error of the mean. DAB = dabrafenib.

### Alteration of iron homeostasis and cellular oxidation in *BRAF^V600E^*-mutant ATC cells by dabrafenib and RSL3 combination therapy

To understand the mechanism of the synergistic anticancer activity of combination dabrafenib and GPX4 inhibitor treatment in *BRAF^V600E^*-mutant ATC cells, we investigated iron homeostasis because ferroptosis is an iron-dependent cell death mechanism (10). We hypothesized that combination therapy altered iron homeostasis, which resulted in cell death (Fig. 3A). In these experiments, all samples were harvested before the onset of cell death. We observed reduced expression of ferroportin-1 (FPN1) in 8505C (Fig. 3B) and SW1736 (Fig. 3C) ATC cell lines following treatment with dabrafenib or non-lethal dose of RSL3. Furthermore, the combination treatment resulted in even greater downregulation of FPN1 compared with the single-agent treatment groups (P < 0.05). As iron homeostasis is regulated by iron storage proteins and cellular importer and exporter of iron, we examined the expression of several major iron modulation–associated proteins (11). We found that ferritin heavy chain 1 (FTH1, a pivotal protein involved in iron storage) was upregulated among all the treatment groups, with the combination treatment group showing higher expression compared with each agent alone, but there was no statistical difference (Fig. 3B, C). Conversely, both divalent metal transporter 1 (DMT1), responsible for free iron uptake into cells, and transferrin receptor 1 (TFR1), which regulates iron uptake from transferrin, were downregulated in the treatment groups (Fig. 3B, C). Next, we measured intracellular ferrous iron and found significant intracellular accumulation in the 8505C (Fig. 3D) and SW1736 (Fig. 3E) ATC cell lines with combination therapy compared with each agent alone (P < 0.001). Taken together, these findings suggested that combination treatment in ATC cells result in iron overload as a result of reduced iron efflux, due to the downregulation of the iron exporter FPN1.

**Figure 3.**
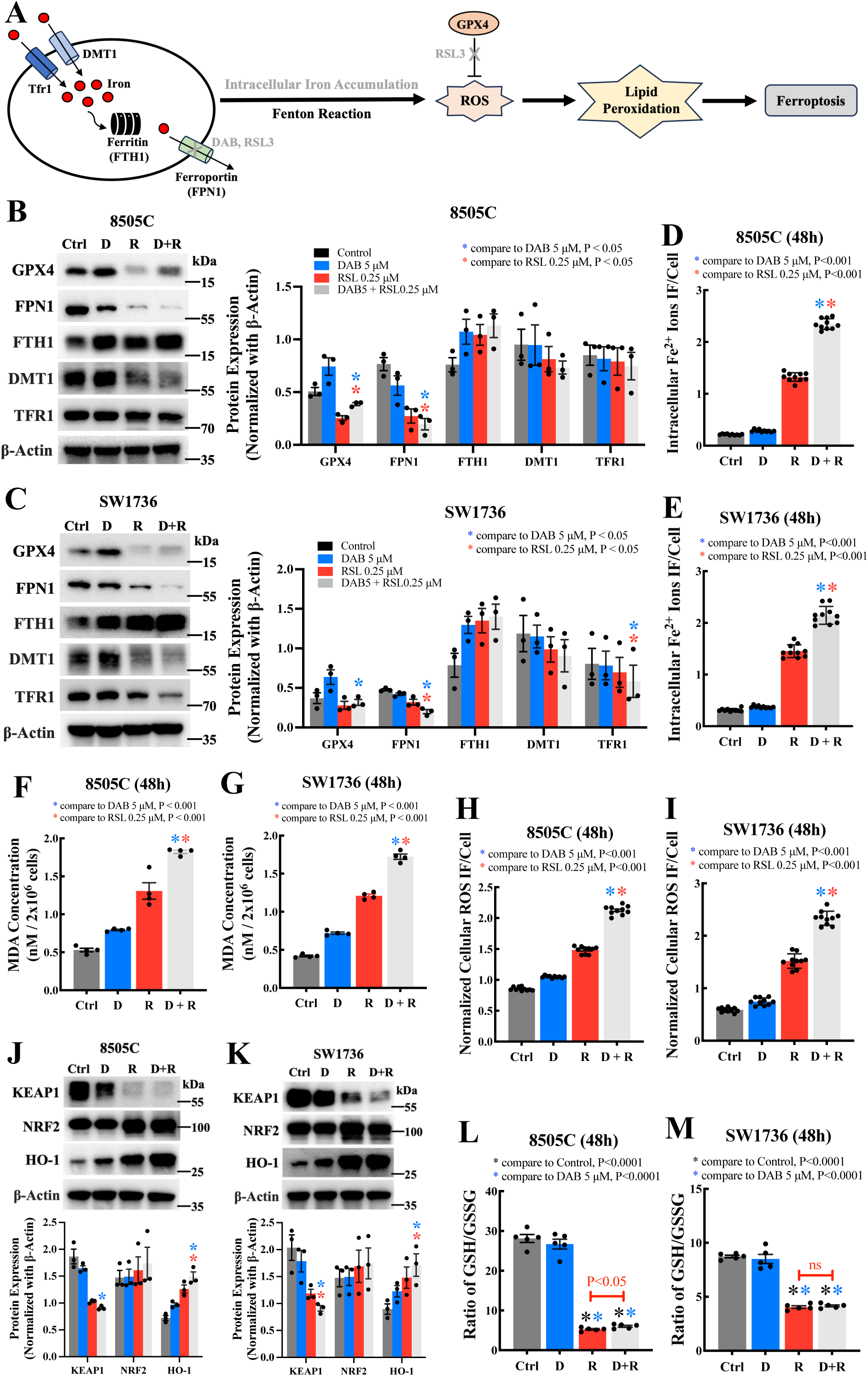
Alterations in iron homeostasis and cellular oxidation in *BRAF^V600E^*-mutant ATC cells with combination dabrafenib and RSL3 treatment. (A) A schematic overview of iron metabolism and ferroptosis. (B, C) Representative images of western blots and analysis of the protein expression levels of GPX4 and proteins related to iron homeostasis. The results are presented as the mean ± the standard error of the mean (n = 3); P values are as indicated in the figure. (D, E) Quantification of intracellular Fe^2+^ by fluorescence intensity (FI) was normalization with the number of total viable cells. The results are presented as the mean ± the standard error of the mean (n = 10); P < 0.001. (F, G) Quantification of the malondialdehyde (MDA) concentration as a measure of lipid peroxidation. This assay was performed using a cell density of 2 × 10^6^ cells per well. The results are presented as the mean ± the standard error of the mean (n = 4); P < 0.001. (H, I) Analysis of cellular reactive oxygen species (ROS) levels after treatment with dabrafenib, RSL3, or their combination for 48 h. The ROS level by fluorescence intensity (FI) was normalized to the number of total viable cells for the quantification. The results are presented as the mean ± the standard error of the mean (n = 10); P < 0.001. (J, K) Representative images of western blots and analysis of protein expression levels of KEAP1–NRF2–HO-1 pathway biomarkers. The results are presented as the mean ± the standard error of the mean (n = 3); P < 0.05. (L, M) The GSH/GSSG ratio in *BRAF^V600E^*-mutant ATC cells after treatments. The results are presented as mean ± the standard error of the mean (n = 5); P values are as indicated in the figure. Ctrl = control, D = dabrafenib at 5 μM, R = RSL3 at 0.25 μM, D+R = dabrafenib (5 μM) in combination with RSL3 (0.25 μM).

Since combination treatment resulted in enhanced intracellular iron overload, we next determined whether it can lead to cellular oxidative stress. We found that lipid perioxidationthe cellular malondialdehyde (MDA) (Fig. 3F, G) and reactive oxygen species (ROS) levels (Fig. 3H, I) were increased significantly compared with each agent alone (P < 0.001). These findings suggest that combination treatment results in synergistic anticancer activity by inducing cellular oxidative stress.

The pathway involving Kelch-like ECH-associated protein 1, Nuclear factor erythroid 2-related factor 2, and Heme oxygenase 1 (KEAP1–NRF2–HO-1) provides an important antioxidant defense mechanism against cellular oxidative stress (12). We found the antioxidant defense mechanism involved in the KEAP1–NRF2–HO-1 pathway was activated in the treatment groups, especially in the combination treatment group, with significant upregulation of HO-1 compared with each agent alone (P < 0.05; Fig. 3J, K).

We also investigated the effect of combination treatment on the canonical pathways of ferroptosis and MAPK signaling. We observed low GPX4 protein level with RSL3 treatment. Interestingly, we found increased GPX4 protein level in the dabrafenib-only and the combination groups as compared with the RSL3-only group (Fig. 3B, C). Similarly, and as expected, phospho-MEK was downregulated in the dabrafenib-only group; however, it was slightly increased in the RSL3-only group compared with the control. The combination treatment group showed rebound increased phospho-MEK level compared with the dabrafenib-only group (Fig. S2). Because GPX4 is a glutathione (GSH)-dependent enzyme, we assessed the functional status of GPX4 after treatments by evaluating the GSH/GSSG ratio. In the dabrafenib-only treatment group, we observed no significant changes compared with the control, while in the RSL3-only and combination treatment groups, we observed a decrease in the GSH/GSSG ratio compared with the control (Fig. 3L, M). Interestingly, we also observed a significant rebound in the 8505C cells (P < 0.05; Fig. 3L) but not in SW1736 cells (nonsignificant; Fig. 3M).

### Combination dabrafenib and RSL3 treatment in tumor spheroids generated from *BRAF^V600E^*-mutant ATC cell lines and a tumor from a patient

Because spheroids derived from ATC cell lines and patient tumors better recapitulate the molecular features of human ATC and response to therapy (13), we used this *ex vivo* experimental model to evaluate the anticancer activity of combination dabrafenib and RSL3 treatment (Fig. 4A). We observed that S-8505C (where S is spheroid), S-SW1736, and patient tumor–derived spheroids (ATC01), all *BRAF^V600E^* mutant, were sensitive to dabrafenib or RSL3 treatment. Furthermore, combination dabrafenib (5 μM) and RSL3 (0.25 μM) treatment significantly reduced cell viability in the *BRAF^V600E^*-mutant ATC spheroids compared with each agent alone (P < 0.001; Fig. 4B–G).

**Figure 4.**
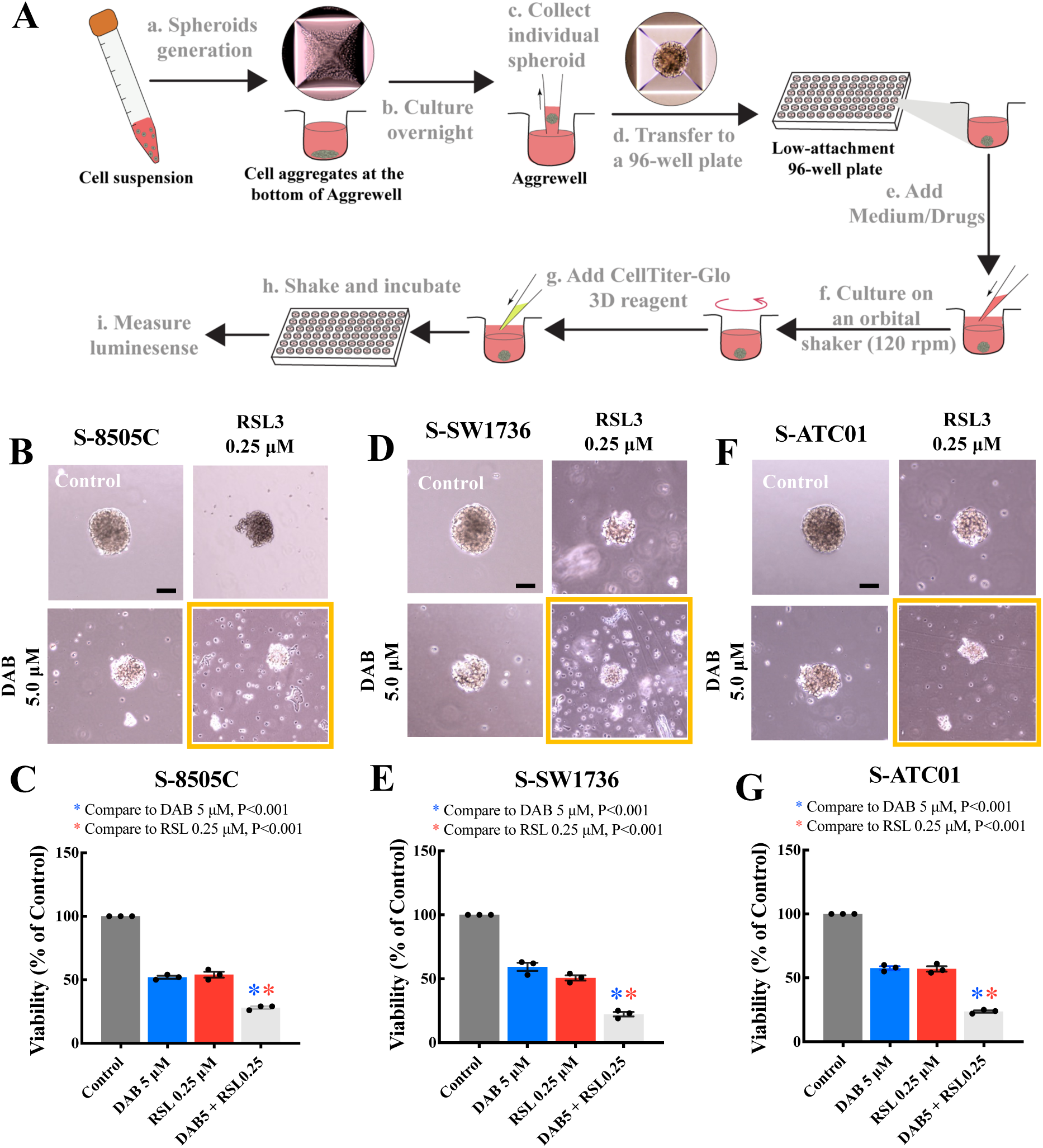
Combination dabrafenib and RSL3 treatment in tumor spheroids generated from the *BRAF^V600E^*-mutant ATC cell lines. (A) The flow diagram shows how the ATC tumor spheroids were generated and the CellTiter-Glo^®^ 3D cell viability assay was conducted. (B, C) Representative images of the 8505C spheroids and the corresponding treatment groups, and cell viability in the 8505C spheroids. (D, E) Representative images of the SW1736 spheroids and the corresponding treatment groups, and cell viability in the SW1736 spheroids. (F, G) Representative images of the ATC01 spheroids and the corresponding treatment groups, and cell viability in the ATC01 spheroids. Cell viability was measured by CellTiter-Glo assay and results were normalized to dimethyl sulfoxide–treated control spheroids. The data are presented as the mean ± the standard error of the mean (n = 3). S = spheroid. The scale bars are 200 μm.

### Combination dabrafenib and GPX4 inhibition (C18) treatment inhibits tumor growth *in vivo*

Next, we tested combination dabrafenib and ferroptosis induction treatment in an *in vivo* model of ATC. Given our *in vitro* findings that targeting GPX4 versus system X_c_^−^ (via erastin) of the ferroptosis pathways with BRAF inhibition have different anticancer activity in *BRAF^V600E^*-mutant ATC cells, we used C18 (a GPX4 inhibitor with *in vivo* activity(14)) and imidazole ketone erastin (IKE, an inhibitor of system X_c_^−^ with *in vivo* activity), respectively, for the *in vivo* studies. Dabrafenib, C18, and IKE treatment alone reduced the tumor volume by 42.4%, 51.4%, and 56.2%, respectively, compared with the vehicle control (Fig. 5A, B). Combination dabrafenib and C18 treatment resulted in a significant reduction in tumor size (76.7%) compared with the vehicle control, dabrafenib alone, and C18 alone (P < 0.001, P < 0.01, and P < 0.05, respectively; Fig. 5A, B).

**Figure 5.**
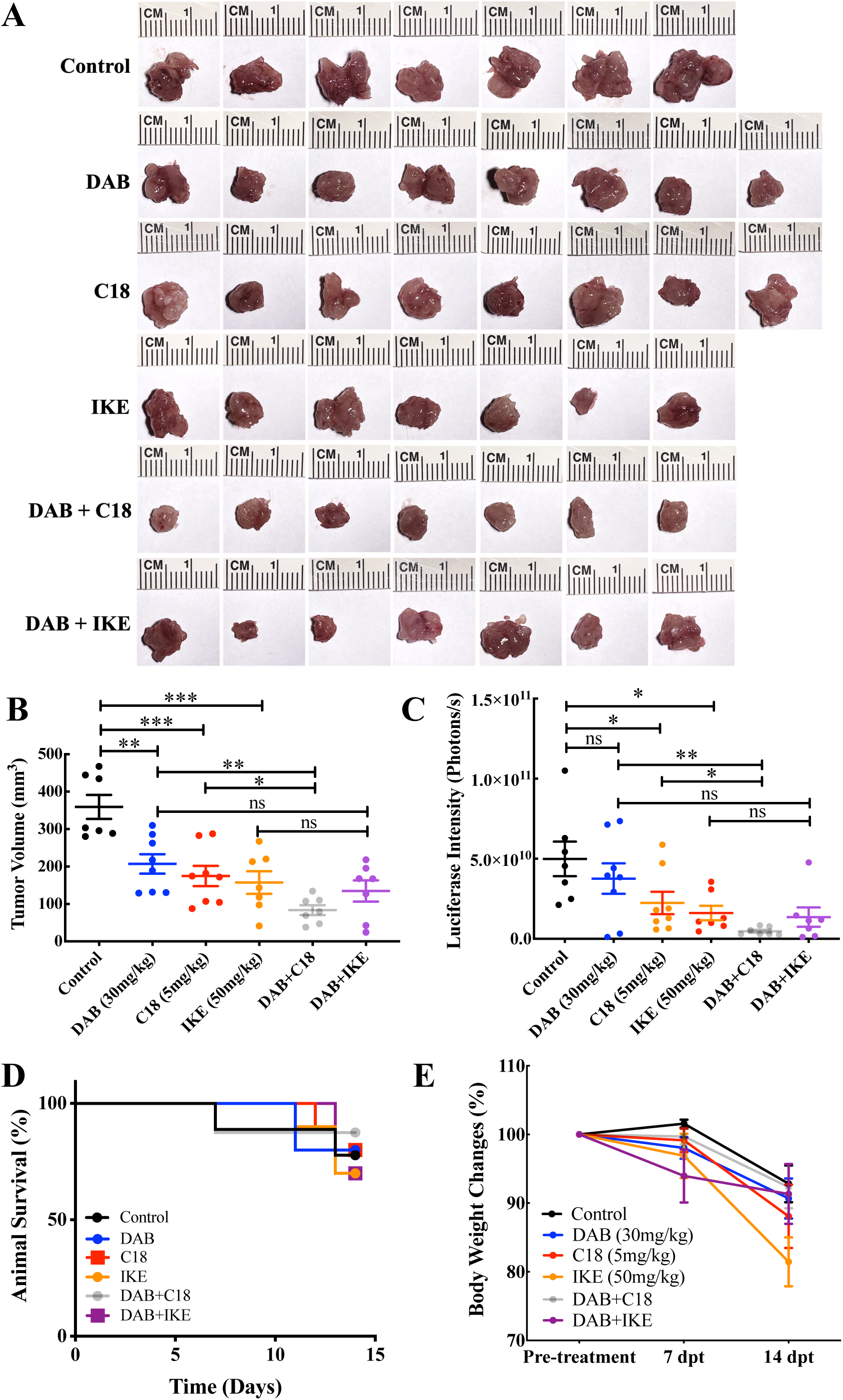
The effects of BRAF inhibition and ferroptosis induction in in the orthotopic xenograft 8505C-Luc2 ATC mouse model. (A) The Kaplan–Meier survival curves of 8505C-Luc2 ATC mice treated with vehicle, single agents, and combinations. (B) Mouse body weight changes 14 days post-treatment. (C) Tumors isolated from treatment groups at 14 days post-treatment. (D) Tumor volume measurements at 14 days post-treatment. (D) Tumor luciferase intensity measurements at 14 days post-treatment. All data are presented as the mean ± the standard error of the mean (n = 7–8). *P < 0.05, **P < 0.01, and ***P < 0.001. DAB = dabrafenib, IKE = imidazole ketone erastin, dpt = days post treatment, ns = nonsignificant.

However, there was no significant difference when comparing combination dabrafenib and IKE treatment with the IKE-only group. Quantification of the bioluminescence tumor luciferase signal intensity correlated well with direct tumor size measurement at 14 days (Fig. 5C).

The Kaplan–Meier survival curve analysis revealed that the groups treated with IKE exhibited a higher incidence of mortality compared with the other groups. Mice receiving dabrafenib in combination with C18 demonstrated the highest survival rate (Fig. 5D). In addition, the group receiving IKE treatment alone reached the humane endpoint at 14 days post-treatment due to notable body weight loss (20% compared with the initial body weight). There were no significant differences in body weight or treatment-related toxicity among the other treatment groups (Fig. 5E). The dabrafenib and C18 combination treatment group exhibited better overall mobility compared with the control and single-agent treatment groups.

### BRAF inhibition–resistant ATC cells are vulnerable to GPX4 inhibition

To test the effects of ferroptosis in BRAF inhibition-resistant *BRAF^V600E^*-mutant ATC cells, we exposed the cells to a high concentration of dabrafenib (40 μM) for more than 3 months. We confirmed that the 8505C and SW1736 ATC cell lines developed resistance to dabrafenib.

Specifically, based on the dose-response curves, the dabrafenib-resistant cells showed significantly lower sensitivity to this drug compared with the corresponding parental cells (P < 0.05; Fig. 6A, B). Importantly, there was increased sensitivity to the GPX4 inhibitor RSL3, indicating that these dabrafenib-resistant cells were more sensitive to ferroptosis induction than their parental cells (P < 0.05; Fig. 6C, D). However, the parental and dabrafenib-resistant *BRAF^V600E^*-mutant ATC cells showed similar sensitivity to erastin (nonsignificant; Fig. 6E, F).

**Figure 6.**
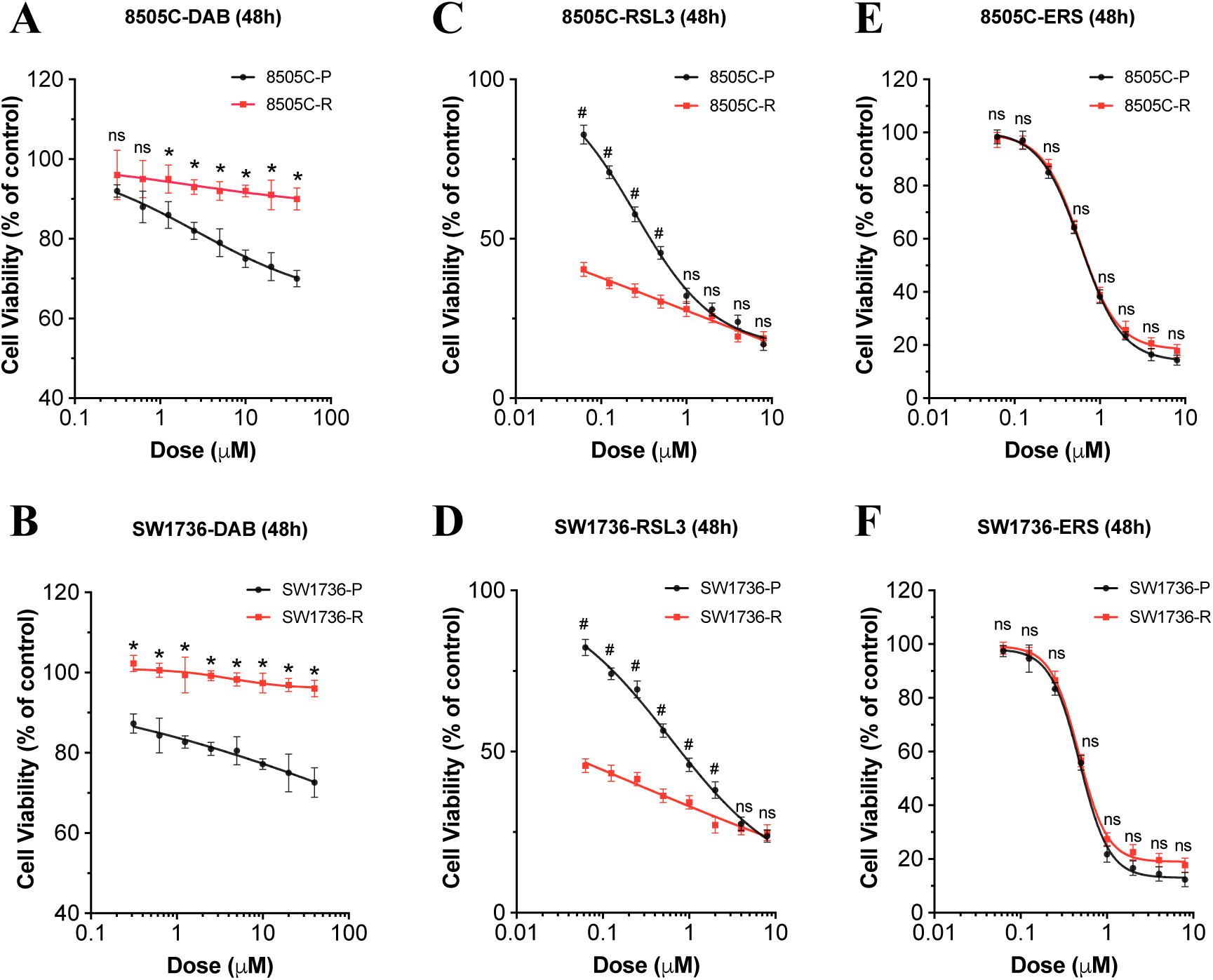
Targeting the ferroptosis pathway in BRAF inhibition–resistant *BRAF^V600E^*-mutant ATC cells. (A, B) Dose-response curves showing decreased sensitivity to dabrafenib in the *BRAF^V600E^*-mutant ATC cell lines with long-term treatment-induced (90 days) dabrafenib-resistance (*P < 0.05). (C, D) Dose-response curves showing increased sensitivity to RSL3 in the dabrafenib-resistant *BRAF^V600E^*-mutant ATC cell lines (^#^P < 0.05). (E, F) Dose-response curves showing equal sensitivity to ferroptosis induction caused by erastin in the parental and resistant cell lines (ns = nonsignificant). The cell viability percentage was calculated relative to the dimethyl sulfoxide– treated control cells. The data are presented as the mean ± the standard error of the mean (n = 3). P = parental, R = resistant, DAB = dabrafenib, ERS = erastin.

## DISCUSSION

In this study, we demonstrated that dual inhibition of BRAF and GPX4 results in synergistic anticancer activity in *BRAF^V600E^*-mutant cells based on *in vitro*, *ex vivo*, and *in vivo* experiments. The synergistic anticancer activity of combination BRAF and GPX4 inhibition is due to increased cellular iron accumulation because of downregulation of the iron exporter FPN1. This, in turn, intensifies the Fenton reaction and oxidative stress, ultimately enhancing ferroptotic cell death. Moreover, we found that dabrafenib-resistant *BRAF^V600E^*-mutant ATC cells are more sensitive to GPX4 inhibition than parental cells, but not to system Xc^−^ inhibition.

Approximately 20% of patients with activating *BRAF* mutations exhibit intrinsic resistance and do not respond to BRAF inhibitors (15–17). Resistance to BRAF inhibition in *BRAF^V600E^*-mutant melanoma cell lines has also been linked to high levels of c-JUN and traits of a mesenchymal-like phenotype (18–20). Therefore, new therapeutic strategies to overcome intrinsic or acquired drug resistance to BRAF inhibition are urgently needed.

There is growing evidence that indicates a mesenchymal or persister cell states significantly relies on GPX4 activity for survival; hence, co-treatment with GPX4 inhibitors may help overcome the resistance mechanisms related to epithelial–mesenchymal transition (EMT) (21–23). In thyroid cancer, GPX4 expression is significantly upregulated compared with normal tissues (23, 24). This overexpression may facilitate thyroid carcinogenesis by inhibiting ferroptosis and is correlated with adverse clinical outcomes (24). In addition, GPX4 acts as a key regulator of ferroptosis, primarily by disrupting lipid peroxidation through the conversion of lipid hydroperoxides into non-toxic lipid alcohols (25). The biosynthesis of GSH and the proper functioning of GPX4 are essential for controlling ferroptosis (26–28). Therefore, dual inhibition of BRAF and GPX4 could exert synergistic anticancer effects in ATC and represent a therapeutic strategy to overcome BRAF inhibitor resistance.

We found that combination GPX4 and BRAF inhibition had synergistic anticancer activity: It inhibited cell proliferation, colony formation, and migration in two *BRAF^V600E^*-mutant ATC cell lines. The molecular mechanisms involved in the synergistic effects of inducting ferroptosis and BRAF inhibition and their downstream influence on cell survival are likely multifactorial. Our study is the first to demonstrate that this novel therapeutic strategy has anticancer activity and mediates intracellular iron overload and cellular oxidative stress and can overcome BRAF inhibitor resistance in *BRAF^V600E^*-mutant ATC cells. Interestingly, GPX4 protein expression rebounded in the combination treatment group relative to the RSL3-only treatment group (Fig. 3B, C). Similarly, the phospho-MEK level was higher in the combination and RSL3-only treatment groups compared with the dabrafenib-only group (Fig. S2). These findings suggest that the observed synergistic anticancer effects are not due to enhanced suppression of the MAPK pathway and induction of ferroptosis with combination therapy. Given iron’s role in generating ROS and facilitating enzyme activity in lipid peroxidation, ferroptosis is tightly regulated by various factors involved in iron metabolism, including iron uptake, storage, utilization, and efflux (29–32). By measuring intracellular free iron, we found that there is a dramatically increased accumulation of iron with combination treatment compared with RSL3 or dabrafenib alone, indicating that iron homeostasis was significantly disrupted. FPN1, also known as SLC40A1, is a transmembrane protein and, at present, is the only putative iron exporter that has been identified (33). It plays a crucial role in regulating systemic iron homeostasis by facilitating the export of cellular iron into the bloodstream (34–36). In addition, studies have reported that activation of the ERK/MAPK pathway can induce FPN1 expression (37), while inhibition of the MAPK signaling pathway can reduce FPN1 expression (11, 38, 39).

Consistent with these previous reports, we also observed a significant reduction in FPN1 protein expression in the dabrafenib- or RSL3-only treatment groups, with an even greater decrease in the combination treatment group. These data strongly support that RSL3 and dabrafenib play a synergistic role in downregulating FPN1, leading to iron accumulation in the cell. This accumulation triggers the Fenton reaction, with consequent overproduction of ROS and buildup of membrane lipid peroxidation, ultimately resulting in ferroptosis.

We found that multiple antioxidative defense mechanisms are involved to cope with the overall cellular stress. For example, at the level of iron metabolism, upregulation of FTH1 and downregulation of DMT1 and TFR1 increased iron storage and reduced iron influx, compensating for the iron accumulation caused by the downregulation of FPN1. In addition, the antioxidant defense mechanism involving the KEAP1–NRF2–HO-1 pathway was activated in all treatment groups, particularly in the combination treatment group. Moreover, there was rebound of GPX4 protein expression and GPX4 activity, as reflected by the GSH/GSSG ratio, in the combination treatment group compared with the RSL3-only treatment group (Fig. 3B–E). This could be due to a cellular compensatory mechanism as the cells attempt to survive the overall cellular stress by increasing GPX4 (22, 40). Similarly, the increased phospho-MEK level in the combination and RSL3-only treatment groups compared with the dabrafenib-only treatment group could also be due to compensatory defense mechanism to overcome the overall increased ROS levels (27, 41). Although the 8505C and SW1736 ATC cell lines are *BRAF^V600E^* mutant, they showed differences regarding the rebound in the protein levels of GPX4 and MAPK pathway components. Compared with the RSL3-only treatment group, GPX4 protein expression and the GSH/GSSG ratio were higher with combination treatment in the 8505C ATC cell line, but not as high in the SW1736 ATC cell line. This could be due to the presence of *TP53* and *PIK3CA* mutations and activation of the PI3K/AKT/mTOR pathway in the 8505C ATC cell line, changes that are closely linked to more aggressive cancer phenotypes (42, 43).

In our previous study, we showed that ATC spheroids closely matched parental patient tumor– derived cells compared with monolayer cultures and exhibit an EMT-associated gene expression profile (13). In addition, the tumor spheroids are typically more resistant to chemo- and radiotherapies than monocultures (13, 44). Therefore, we tested the effects of the combination treatment on *ex vivo* spheroids. We found synergistic anticancer activity in the combination group in this three-dimensional disease model, indicating the strong potential of the combination therapeutic effects in human ATC. Due to the poor bioavailability of RSL3 *in vivo* (14, 45), we used C18, a recently reported biocompatible GPX4 inhibitor, in the *in vivo* study (14). Combination C18 and dabrafenib treatment exhibited prominent antitumor activity *in vivo* compared with other groups.

The differences between erastin and RSL3 sensitivity in the parental and dabrafenib-resistant *BRAF^V600E^*-mutant ATC cells could be attributed to their upstream and downstream targets, respectively, of the ferroptosis pathway. Factors such as those affecting the ability of erastin to inhibit the upstream system Xc^−^ target or the ability of cells to compensate for cystine/glutathione depletion could reverse erastin sensitivity without modulating RSL3 sensitivity.

In summary, we demonstrated that dual targeting of ferroptosis (GPX4) and *BRAF^V600E^* has significant and synergistic anticancer activity in *BRAF^V600E^*-mutant ATC cells *in vitro*, *ex vivo*, and *in vivo*. This synergistic activity is mediated by enhanced cellular oxidative stress due to iron overload. Combination GPX4 inhibition and dabrafenib treatment is a promising treatment strategy for *BRAF^V600E^*-mutant cancers.

## MATERIALS AND METHODS

### Chemistry

All the chemical agents and solvents were purchased from commercial suppliers. Dabrafenib (S2807), RSL3 (S8155), ML162 (S4452), erastin (S7242), IKE (S8877), ferrostatin-1 (S7243), and PEG300 (50-217-3211) were purchased from Selleckchem (Houston, TX, USA). GPX4-IN-5 (compound C18) was purchased from MedChemExpress (Monmouth Junction, NJ, USA). Dimethyl sulfoxide (DMSO, J66650) was purchased from Thermo Fisher Scientific (Rockford, IL, USA). Tween 80 (ICN10317080) was purchased from MP Biomedicals (Santa Ana, CA, USA).

### Cell lines and culture conditions

The ATC cell lines 8505C and SW1736, harboring the *BRAF^V600E^* mutation, were purchased from the European Collection of Cell Culture (Salisbury, United Kingdom) and Cell Lines Service GmbH (Eppelheim, Germany), respectively. The *BRAF*^WT^ cell line THJ16T derived from a patient with ATC was kindly gifted by Dr. John A. Copland (Mayo Clinic, Jacksonville, FL, USA). Patient tumor–derived ATC01 spheroids were generated and established in our laboratory as reported previously (13). All cell lines used were maintained in Dulbecco’s Modified Eagle’s Medium (DMEM, which contains 4,500 mg/L of D-glucose, 2 mmol/L of L-glutamine, and 110 mg/L of sodium pyruvate) supplemented with 10% fetal bovine serum (FBS), penicillin (10,000 U/mL), streptomycin (10,000 U/mL), and fungizone (250 ng/mL) to make the complete medium. Cell cultures were maintained at 37°C with a 5% CO_2_ and 95% O_2_ atmosphere within a standard humidified incubator.

### Cell viability assay

Cell viability assays were performed in triplicate to evaluate the effect of the drugs on cell viability. Cells were plated in 96-well black-bottom plates (Greiner Bio-One, Kremsmünster, Austria) at 1.5 × 10^3^ cells/well in 100 μL of complete culture medium. After 24 h, adhered cells were treated with different doses of dabrafenib, RSL3, ML162, and erastin (Selleckchem), alone or in combination in 100 μL of fresh complete medium. DMSO was used as the vehicle. Plates were collected at 48 h after treatment, and the cell viability assay was performed using the CyQUANT kit according to manufacturer’s instructions (Thermo Fisher Scientific, C7026). The number of cells was determined by using a fluorescence microplate reader (Molecular Devices, San Jose, CA, USA) at 485 nm/538 nm; the results are expressed in relative fluorescence units (RFU). The CI of cell viability suppression induced by the combination treatment was calculated using the Chou–Talalay method. A value < 1 indicates a synergistic effect; the lower the value, the stronger the synergistic effect (46).

### Clonogenic assay

Cells were plated in 12-well plates (5 × 10^2^ cells/well) and allowed to adhere overnight in complete culture medium. Then, they were treated with different doses of dabrafenib or RSL3 alone or in combination for 24 h (for short-term treatment), followed by washing and the addition of fresh complete medium. The colonies were allowed to grow for 12–14 days. For long-term treatment, the cells were treated for 12 days. The growth medium was replaced every 72 h. The colonies were fixed with 0.4% buffered paraformaldehyde and then stained with 0.5% crystal violet in methanol for 10 minutes. The colonies were photographed and counted by using the ChemiDoc Imaging System (Bio-Rad, Hercules, CA, USA) and the ImageJ software (National Institutes of Health, Bethesda, MD, USA).

### Migration assay

A scratch plate or wound healing assay was performed to assess the effect of the drugs on cell migration. First, 1.5 × 10^5^ cells were plated in 6-well plates and allowed to attach and grow for 24 h. Then, they were treated with vehicle, dabrafenib (5 μM), or RSL3 (0.25 μM) alone or in combination for 24 h. The concentrations used in this assay have minimal effects on cell viability alone after 24 h treatment, ensuring that the results accurately reflect cell migratory behavior. A cross-pattern wound was made with a sterile 200-µL pipette tip. The cells were washed with phosphate-buffered saline (PBS), and then fresh medium with 2% serum was added. Images were captured every 8 h and measurements were made up to 24 h. The ImageJ software was used to measure the wound area. The experiments were performed in triplicate.

### Lipid peroxidation assay

*BRAF^V600E^*-mutant ATC cells were plated in a 10-cm culture dish overnight. Subsequently, the medium containing vehicle, RSL3 (0.25 μM), dabrafenib (5 μM), or their combination was added and incubated for 48 h. Following treatment, the culture medium was aspirated, and the cells were washed with PBS. Subsequently, 2 × 10^6^ cells from each group were collected and lysed using a Dounce homogenizer on ice. The lysates were then centrifuged at 13,000 *g* at 4°C for 10 minutes to obtain the supernatant. The concentration of malondialdehyde (MDA) was determined as per the manufacturer’s instructions using the Lipid Peroxidation Assay Kit (Abcam, Milpitas, CA, USA, ab118970).

### Cellular ROS assay

Intercellular ROS formation was assessed using the DCFDA/H2DCFDA-Cellular ROS Assay Kit (Abcam, ab113851) and performed according to the manufacturer’s instructions. The results were normalized based on the number of cells obtained from parallel cell proliferation plates.

### Determination of the GSH/GSSG ratio

*BRAF^V600E^*-mutant ATC cells were seeded in 96-well black-bottom plates (Greiner Bio-One) at 1 × 10^4^ cells/well in 100 μL of complete culture medium. Subsequently, the medium containing either vehicle, RSL3 (0.25 μM), dabrafenib (5 μM), or their combination was added and incubated for 48 h. After treatment, the levels of GSH and GSSG were assessed using a commercial GSH/GSSG assay kit (Promega, Madison, Wisconsin, USA, V6611), according to the manufacturer’s protocol.

### FerroOrange assay

ATC cells were seeded in 96-well black-bottom plates (Greiner Bio-One) at a density of 1.5 × 10^3^ cells per well in 100 μL of complete culture medium. Subsequently, the medium containing vehicle, RSL3 (0.25 μM), dabrafenib (5 μM), or their combination was added and incubated for 48 h. Upon completion of the treatment period, the culture medium was aspirated, and the cells were washed with serum-free medium. Subsequently, the cells were incubated with the FerroOrange probe for 30 minutes at 37°C to facilitate the binding of labile iron ions within the cells. Excess FerroOrange probe was removed by washing with serum-free medium. The fluorescence intensity of the bound FerroOrange was measured using a microplate reader, with subsequent normalization based on the number of cells obtained from parallel cell proliferation plates.

### Western blot analysis

Total cell lysates were prepared from the ATC cells after treatment with drugs alone or in combination or vehicle using radioimmunoprecipitation assay (RIPA) buffer (Thermo Fisher Scientific) and 1% sodium dodecyl sulfate (SDS). The protein concentration was determined using the Pierce™ BCA protein assay kit (Thermo Fisher Scientific). Equal amounts of proteins were resolved by electrophoresis on 10%–15% SDS-polyacrylamide gel electrophoresis (PAGE) gels, transferred to a nitrocellulose membrane (Bio-Rad), and incubated overnight with specific primary antibodies. On the next day, unbound primary antibody was washed away, and the membrane was incubated with secondary antibody. The blots were then developed using an ECL substrate. The blots were visualized with the ChemiDoc Imaging System. The ImageJ software was used to quantify protein bands with densitometry. Table S2 provides detailed information on the antibodies used.

### Tumor spheroid generation and drug testing

According to a published study (13), tumor spheroids derived from *BRAF^V600E^*-mutant ATC cells were generated in a 24-well AggreWell plate (STEMCELL Technologies, Vancouver, British Columbia, Canada, cat#34411). Tumor spheroids were formed after centrifugation (350 *g*, 5 minutes), and the plate was incubated overnight (37°C, 5% CO_2_). On day 2, individual tumor spheroids were transferred to low-attachment 96-well plates under a microscope; each well contained 50 μL of medium. Next, 50 µL of the appropriate treatment was added to each (the control group was DMSO). Then, the plates were placed on an orbital shaker (120 rpm) to provide a floating environment and to ensure adequate interaction with the drugs. Redosing was performed after 3 days, and viability of the spheroids was determined after 5 days of treatment. After treatment, 100 μL of CellTiter-Glo 3D reagent (Promega, cat#G9681) was added to each well. After shaking and incubation, the luminescence was measured and recorded.

### Orthotopic xenograft ATC models

Experimental protocols were approved by the Stanford University Administrative Panel on Laboratory Animal Care (APLAC). For orthotopic implantation of ATC cells, 1 × 10^6^ 8505C-Luc2 cells (where Luc2 indicates cells with stable expression of a luciferase reporter) were surgically implanted into the ride side region of the thyroid gland in *NOD Cg-Prkdcscid Il2rgtm1WjI/SzJ* mice. Both male and female mice were used in this study. Tumor luminescence was monitored using the IVIS *in vivo* imaging system (Xenogen, Alameda, CA, USA) following intraperitoneal injection of luciferin (10 μg/kg body weight). One week after orthotopic implantation, the mice were randomly assigned into six treatment groups and started with treatments. Dabrafenib was dissolved in 0.5% methylcellulose fiber in ddH_2_O under stirring at 4°C until completely dissolved. A bioavailable GPX4 inhibitor, compound C18, was prepared in solvent (DMSO/PEG300/Tween80/saline = 5%:40%:5%:50%) (14). Group I control mice received vehicle control via oral gavage and an intraperitoneal (IP) injection of blank solvent daily. Group II mice received dabrafenib (30 mg/kg body weight) via oral gavage along with an IP injection of blank solvent daily. Group III mice received vehicle control via oral gavage and an IP injection of C18 (5 mg/kg body weight) daily. Group IV mice received vehicle control via oral gavage and an IP injection of IKE (50 mg/kg body weight) daily. Group V mice received dabrafenib (30 mg/kg weight) via oral gavage and an IP injection of C18 (5 mg/kg body weight) daily. Group VI mice received dabrafenib (30 mg/kg body weight) via oral gavage and an IP injection of IKE (50 mg/kg body weight) daily. Mice underwent weekly imaging and weight measurements throughout the 14-day treatment period, after which they were euthanized via CO_2_ inhalation. Tumor, lungs, and liver tissues were collected post-euthanasia for histological analysis.

### Statistical analyses

The data are presented as the mean ± standard error of the mean or the mean ± standard deviation, and the details for each analysis are provided in the figure legends. For parametric data, analysis of variance (ANOVA), with appropriate *post hoc* tests, was used to determine the differences between the groups. A two-tailed P value of ≤ 0.05 was considered to be statistically significant. GraphPad Prism version 8 (GraphPad Software Inc., San Diego, CA, USA) was used to perform all statistical analyses. The level of statistical significance is denoted in the figures and figure legends.

## Acknowledgments

We would like to acknowledge the experimental support from the Stanford Animal Care and Facilities for helping with the animal study.

## Authors’ contributions

EK, SJD, and JH contributed to the study concept. JH, CG, TP, ZY, and YT contributed to the *in vitro* and *ex vivo* data collection and analysis. JH, CG, TP, ZY, and EDA contributed to the data collection and analysis for the animal studies. JH, CG, TP, ZY, MB and EK contributed to the manuscript preparation. EK supervised the study and coordinated the research efforts and writing of the manuscript.

## Data and material availability

The datasets generated and analyzed during the current study are available on reasonable request.

## List of Supplementary Materials

**Supplementary Fig S1.**
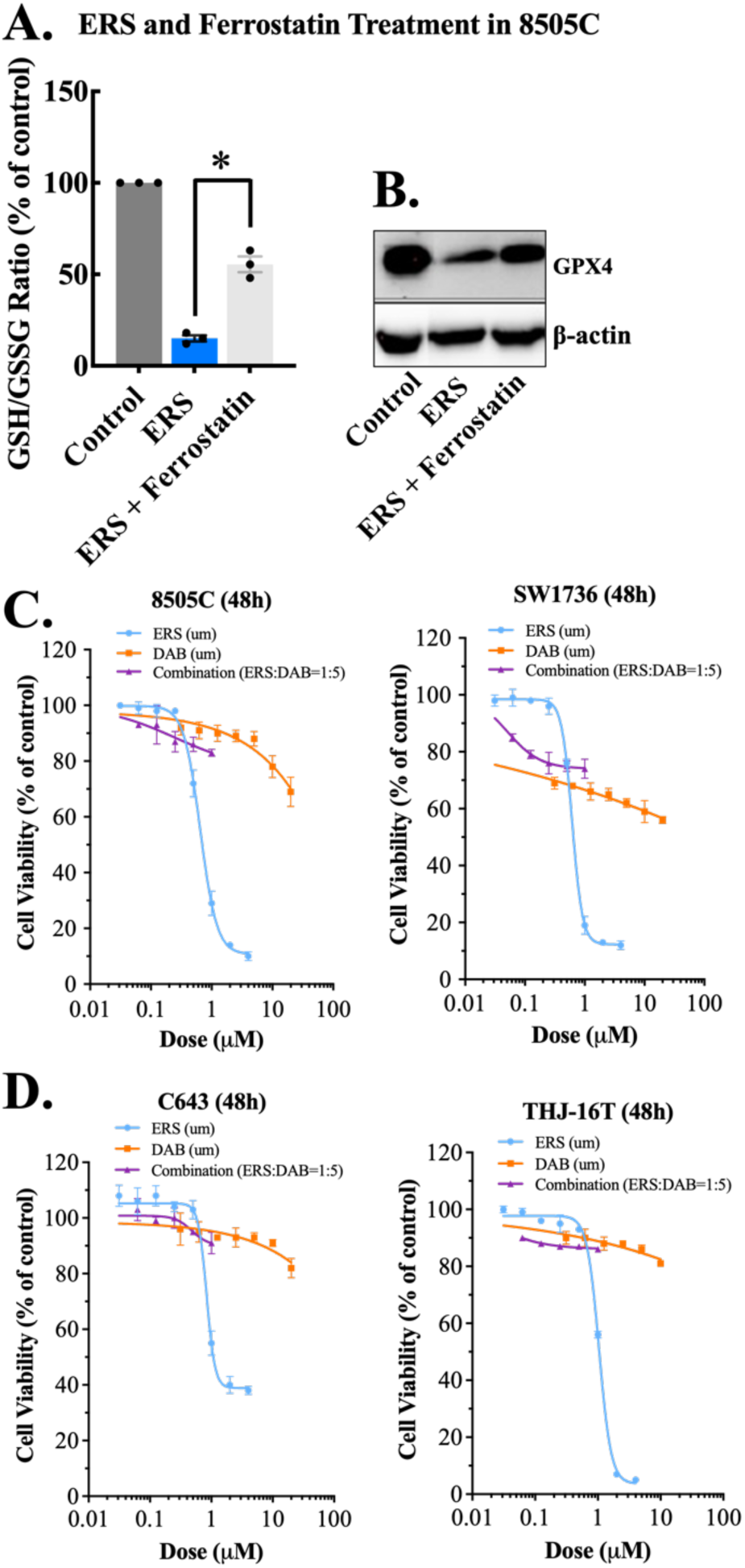
Erastin treatment in ATC cells. (A) The GSH/GSSG ratio and (B) expression level of GPX4 were reduced with erastin (2 μM) in a *BRAF*^V600E^-mutant 8505C ATC cell line for 48h, which was partially recovered by ferrostatin-1 (a ferroptosis inhibitor, 2 μM), confirming induction of ferroptosis in the ATC cells. (C) The cell viability curves of combinatorial treatment in *BRAF^V600E^*-mutant (C) and *BRAF^WT^* (D) ATC cell lines were plotted according to the doses of RSL3 used in the treatment. *p < 0.05, ns = nonsignificant. All data were presented as mean ± SEM (n = 3).

**Supplementary Fig S2.**
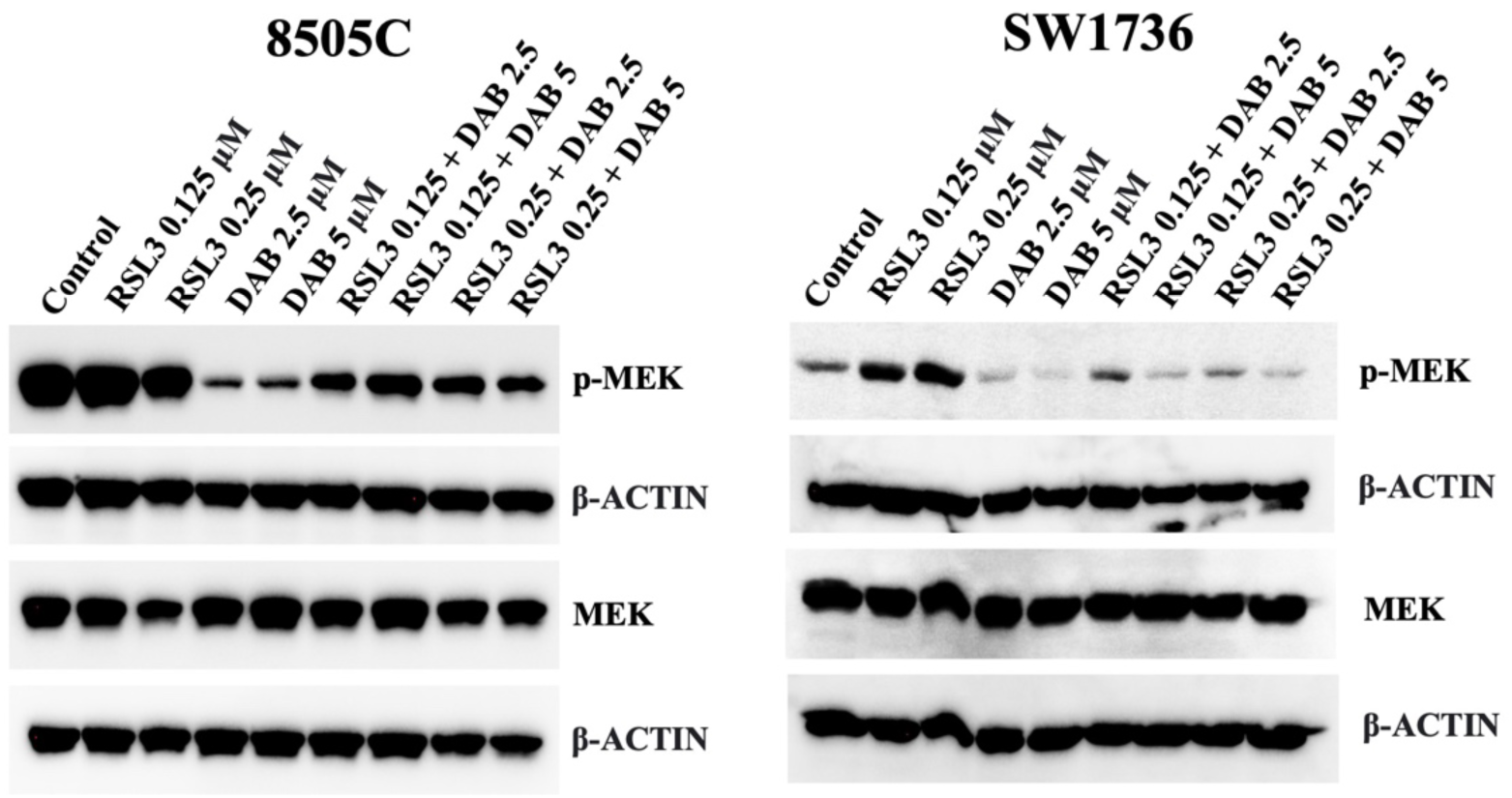
Effect of dabrafenib and RSL3 combination on MEK pathway.

**Supplemental Table S1.** Dabrafenib (D) and RSL3 (R) combination shows synergistic activity on *BRAF^V600E^*-mutant thyroid cancer cells.

| Cell lines | D10 +<br>R0.5<br>μM | D10 +<br>R0.25<br>μM | D10 +<br>R0.125<br>μM | D5 +<br>R0.5<br>μM | D5 +<br>R0.25<br>μM | D5 +<br>R0.125<br>μM | D2.5 +<br>R0.25<br>μM | D2.5 +<br>R0.125<br>μM |
| --- | --- | --- | --- | --- | --- | --- | --- | --- |
| 8505C | 1.0056 | 1.0901 | **0.4637** | 1.5798 | **0.5336** | **0.3518** | **0.2694** | **0.2018** |
| SW1736 | 3.0365 | 2.3083 | 1.8999 | **0.3299** | **0.4100** | **0.3700** | **0.4700** | **0.3700** |
| C643 | 4.3423 | 1.8103 | 1.0835 | 3.0745 | 2.1360 | 1.4322 | 1.0653 | 1.2184 |
| THJ-16T | 1.6498 | 1.1602 | 1.0063 | 2.1657 | 4.5079 | 5.7678 | 15.516 | 9.8255 |

The combination index (CI) was calculated using the Chou–Talalay method with the combination dabrafenib and RSL3 treatment at 48 h. Dabrafenib (5 μM, 2.5 μM) in combination with RSL3 (0.25μM, 0.125μM) showed synergistic effects in the *BRAF*<sup>V600E</sup>-mutant cell lines (8505C and SW1736). In the *BRAF*<sup>WT</sup> cell lines (C643 and THJ-16T), however, there were mostly antagonistic effects in the combination treatment. The CI was interpreted as follows: < 1, synergist (bold); 1, additive; > 1, antagonist.

**Supplemental Table S2.**
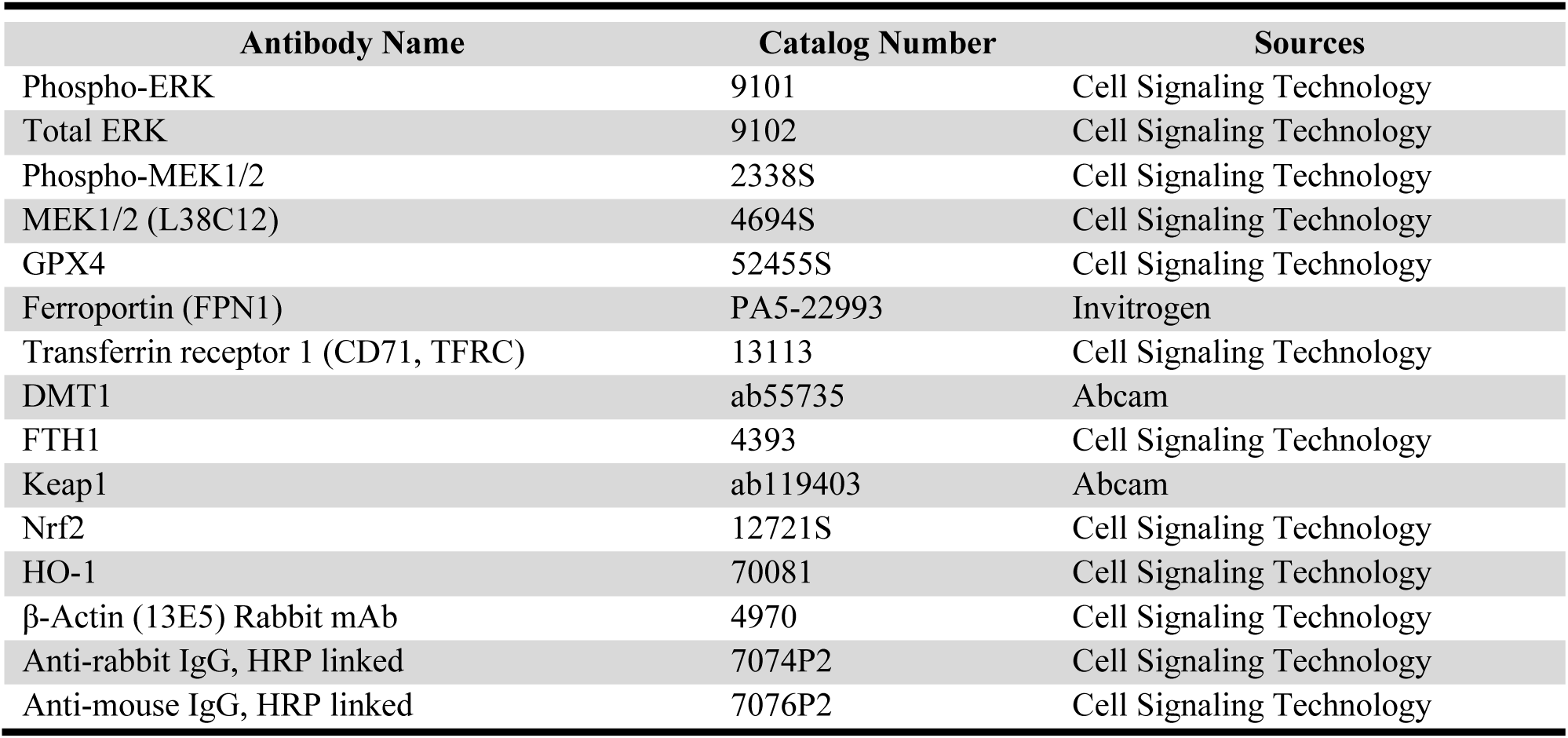
The list of primary and secondary antibodies used in the immunoblotting study.

